# Extracellular S100B alters spontaneous Ca^2+^ fluxes in dopaminergic neurons via L-type voltage gated calcium channels: Implications for Parkinson’s disease

**DOI:** 10.1101/2021.02.24.432751

**Authors:** Eric A. Bancroft, Gauri Pandey, Sara M. Zarate, Rahul Srinivasan

## Abstract

Parkinson’s disease (PD) is associated with an abnormal increase in S100B within the midbrain and cerebrospinal fluid. In addition, overexpression of S100B in mice accelerates the loss of substantia nigra pars compacta (SNc) dopaminergic (DA) neurons, suggesting a role for this protein in PD pathogenesis. We found that in the mouse SNc, S100B labeled astrocytic processes completely envelop the somata of tyrosine hydroxylase (TH) positive DA neurons. Based on this finding, we rationalized that abnormal increases in extracellularly secreted S100B by astrocytic processes in the SNc could alter DA neuron activity, thereby causing dysregulated midbrain function. To test this hypothesis, we measured the effect of bath perfused S100B peptide on the frequency and amplitude of spontaneous calcium fluxes in identified TH^+^ and TH^−^ midbrain neurons from 3-week old mouse primary midbrain cultures. Acute exposure to 50 pM S100B caused a 2-fold increase in calcium flux frequency only in TH^+^ DA neurons. The L-type voltage gated calcium channel (VGCC) inhibitor, diltiazem eliminated S100B-mediated increases in DA neuron calcium flux frequency, while the T-type specific VGCC blocker mibefradil failed to inhibit the stimulatory effect of S100B. Chronic exposure to S100B caused a 3-fold reduction in calcium flux frequencies of TH^+^ neurons and also reduced calcium flux amplitudes in TH^−^ neurons by ∼4-fold. Together, our results suggest that exposure to S100B pathologically alters spontaneous calcium activity in midbrain neurons via an extracellular mechanism involving L-type VGCCs expressed in DA neurons. These findings are relevant to understanding mechanisms underlying DA neuron loss during PD.

**Table of Contents Image:** 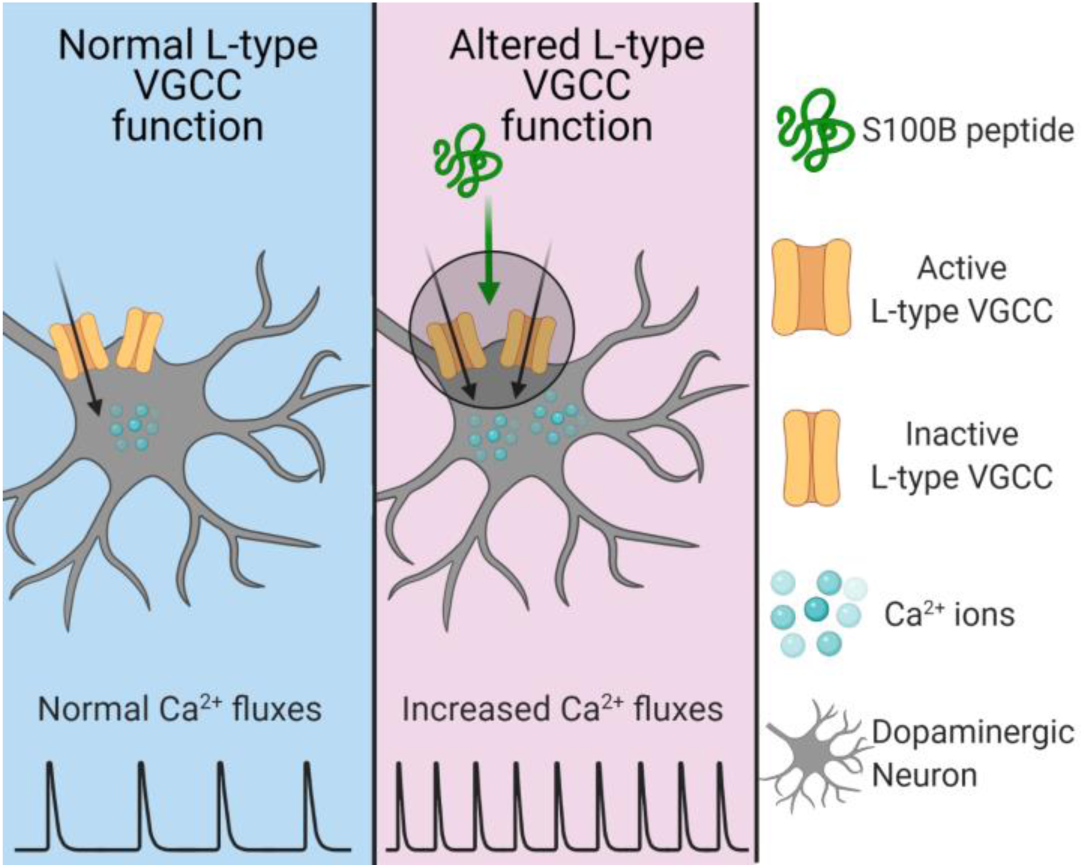

**Main Points:**

- Extracellular S100B increases Ca^2+^ fluxes in dopaminergic neurons
- L-type VGCCs in dopaminergic neurons are required for S100B-mediated increases in Ca^2+^ fluxes
- Chronic S100B alters Ca^2+^ fluxes in dopaminergic and non-dopaminergic neurons

## Introduction

Parkinson’s disease (PD) is the second most common neurodegenerative disorder, projected to reach pandemic proportions by 2040 (Dorsey, Sherer, Okun, & Bloem, 2018). Understanding the mechanisms by which dopaminergic (DA) neurons degenerate in PD is therefore vital for developing effective neuroprotective treatments that can slow or even stop the progressive loss of DA neurons.

Interestingly, patients with PD display increased levels of S100B in their cerebrospinal fluid (CSF) and serum (Sathe et al., 2012; Schaf et al., 2005), as well as significantly higher S100B expression in the substantia nigra pars compacta (SNc) (Sathe et al., 2012). Several lines of evidence show that apart from being merely associated with PD, upregulated S100B can also actively contribute to the degeneration of DA neurons. Specifically, studies show that: (**i**) A single nucleotide polymorphism, rs9722, which is associated with increased levels of serum S100B also results in an elevated risk for early onset PD (Fardell et al., 2018; Hohoff et al., 2010), (**ii**) Ablation of S100B in mice protects against MPTP-induced DA degeneration (Sathe et al., 2012), (**iii**) Mice overexpressing S100B develop parkinsonian features (Liu et al., 2011; Sathe et al., 2012), and (**iv**) Overnight S100B elevation correlates with increased PD severity and sleep disruption (D. Z. Carvalho et al., 2015). Taken together, these reports converge on the idea that abnormal levels of S100B in the SNc could trigger the degeneration of DA neurons in PD patients.

During PD, midbrain astrocytes become reactive and demonstrate a pathological increase in the expression levels of astrocyte-specific proteins such as glial fibrillary acid protein (GFAP) (Batassini et al., 2015; Thannickal, Lai, & Siegel, 2007) and S100B (Sathe et al., 2012). Among the many proteins that are upregulated in reactive astrocytes, S100B is of particular interest in the context of PD because of its potential to initiate pathological processes in DA neurons during early stage PD. Indeed, extracellular S100B has been shown to alter neuronal activity in multiple brain regions (Morquette et al., 2015; Ryczko et al., 2021), and extracellularly secreted S100B from astrocytes accelerates neurodegeneration by engaging receptor for advanced glycation endproducts (RAGE)-mediated pro-inflammatory pathways on astrocytes and microglia (Hofmann et al., 1999; Huttunen et al., 2000; Riuzzi, Sorci, Beccafico, & Donato, 2012). In addition to signaling through RAGE receptors, S100B and the S100 family of proteins interact intracellularly with several ion channels and receptors expressed in neurons, and these interactions result in significant biological effects such as increased neurotropism or the modulation of neuronal excitability (Hermann, Donato, Weiger, & Chazin, 2012).

In this study, we show that acute extracellular exposure to 50 pM of S100B peptide specifically increases the frequency of spontaneous Ca^2+^ fluxes in tyrosine hydroxylase (TH) expressing cultured mouse midbrain neurons via a functional interaction with L-type VGCCs. We also demonstrate that chronic exposure of primary mouse midbrain cultures to extracellular S100B peptide causes a dysregulation of spontaneous Ca^2+^ fluxes in both TH^+^ and TH^−^ midbrain neurons. Given the important roles of L-type VGCCs (Ilijic, Guzman, & Surmeier, 2011; Olson et al., 2005; Sun et al., 2017), astrocytes (Booth, Hirst, & Wade-Martins, 2017; Gomez et al., 2019; Lang et al., 2019) and S100B in the midbrain and PD, these results could have important implications for understanding mechanisms by which an extracellularly secreted astrocytic protein such as S100B can cause dysregulated VGCC function in the midbrain during early stages of PD.

## Materials and Methods

### Mice

All experimental procedures with mice were approved by the Texas A&M University Institutional Animal Care and Use Committee (IACUC), animal use protocol numbers 2017-0053 and 2019-0346. Adult (3-4 month) male C57BL/6 wildtype mice and embryos (ED14) from timed-pregnant female C57BL/6 mice were obtained from the Texas A&M Institute for Genomic Medicine (TIGM). Pregnant mice were group housed in a temperature controlled environment on a 12:12 h light:dark cycle with food and water available *ad libitum*.

### Immunostaining of mouse brain sections and DA cultures

Mice were transcardially perfused with phosphate-buffered saline (PBS, ThermoFisher, Waltham, MA, USA) followed by 10% Formalin/PBS (VWR, Radnor, PA, USA). Brains were postfixed in 10% Formalin/PBS for 24-48 h at 4°C, then moved to 30% sucrose (Sigma-Aldrich, St. Louis, MO, USA) in PBS for dehydration. Brains were sectioned coronally at 40 μm on a sliding microtome (SM2010 R, Leica, Nussloch, Germany). Brain sections were permeabilized in 0.5% Triton X-100 (Sigma-Aldrich) in PBS, and blocked in 10% normal goat serum (NGS, Abcam, Cambridge, UK) in PBS. Primary antibodies used were rabbit polyclonal anti-GFAP (1:1000, Abcam), rabbit polyclonal anti-S100 (1:1000, Abcam) and chicken polyclonal anti-TH (1:1000, Abcam). Secondary antibodies used were goat polyclonal anti-rabbit Alexa Fluor 488 (1:1000, Abcam) and goat polyclonal anti-chicken Alexa Fluor 594 (1:1000, Abcam). Antibodies were incubated in a 1% NGS/PBS solution. For each midbrain subregion, 2 fields of view (FOV) from 2-3 sections per mouse (3 mice total) were used to quantify S100B and TH protein labeling.

DA cultures were fixed by placing coverslips in 10% Formaldehyde/PBS for 40 min. Cultures were permeabilized in 0.01% Triton X-100/PBS and blocked in 10% NGS/PBS solution. Antibodies used in sections were also used in cultures, with the addition of mouse polyclonal anti-NeuN (1:1000, Abcam) and goat polyclonal anti-mouse Alexa Fluor 405 (1:1000, Abcam). Imaging was performed using a confocal microscope (Fluoview 1200, Olympus, Tokyo, Japan) with a 60X and 1.35 NA oil-immersion objective (Olympus).

### Culturing primary mouse midbrain neurons

For culturing primary mouse midbrain neurons, neurobasal medium, DMEM + GlutaMAX medium, GlutaMAX supplement, B-27, equine serum, and penicillin-streptomycin were purchased from ThermoFisher. Deoxyribonuclease I (DNase), poly-L-lysine, poly-L-ornithine, laminin, ascorbic acid, kanamycin, and ampicillin were purchased from Sigma-Aldrich. Corning 35 mm uncoated plastic cell culture dishes were purchased from VWR, 12 mm circular cover glass No. 1 was purchased from Phenix Research Products (Candler, NC, USA). Papain was purchased from Worthington Biomedical Corporation (Lakewood, NJ, USA).

Detailed methods to culture primary mouse DA neurons have been previously described (Bancroft & Srinivasan, 2020; Henley et al., 2017; Zarate et al., 2020). Briefly, cultures were obtained from embryonic day (ED14) mouse embryos of mixed sexes. Timed-pregnant mice (Texas A&M Institute for Genomic Medicine) were sacrificed via cervical dislocation and embryos were removed. Embryos were decapitated and ventral midbrain was dissected using previously described methods (Bancroft & Srinivasan, 2020; Henley et al., 2017). Following dissection, cells were digested in papain for 15 min at 37°C, then cells were separated using DNase treatment and mechanical trituration in a stop solution of 10% equine serum in PBS. Cells were plated at a density between 200,000-300,000 cells per cover glass on 12 mm circular cover glasses triple coated with poly-L-lysine, poly-L-ornithine and laminin. After plating, cells were placed in an incubator at 37°C with 5% CO_2_ for 1 h, followed by addition of 3 ml of neurobasal media supplemented with GlutaMAX, B-27, equine serum, ascorbic acid and containing penicillin-streptomycin, kanamycin, and ampicillin. Culture medium was exchanged at 3 d intervals and all primary mouse midbrain neuron-astrocyte co-cultures were maintained for at least 3 weeks before performing experiments.

### Adeno-associated virus (AAV) vectors

In order to image spontaneous Ca^2+^ fluxes in visually identified DA neurons from primary mouse midbrain cultures, we utilized adeno-associated viruses (AAVs) that report TH^+^ DA neurons (AAV 2/5 TH-tdTomato), while simultaneously allowing the measurement of spontaneous Ca^2+^ fluxes in TH^+^ and TH^−^ midbrain neurons (AAV 2/5 hSyn-GCaMP6f). AAV 2/5 hSyn-GCaMP6f was purchased from Addgene (Cat # 100837-AAV, RRID: Addgene_100837). AAV 2/5 TH-tdTomato was packaged by Vector Builder (Chicago, IL). The targeting plasmid for packaging AAV 2/5 TH-tdTomato was cloned from a pAAV-mouse THp-eGFP plasmid obtained as a gift from Dr. Viviana Gradinaru (California Institute of Technology, Pasadena, CA). The eGFP cassette in the pAAV-mouse THp-eGFP plasmid was replaced with tdTomato between restriction sites Not1 and BamH1

All AAV infections were performed at 14 days in vitro (DIV), as previously described (Bancroft & Srinivasan, 2020). For AAV infections, the culture medium was removed and 1 ml of serum-free DMEM + GluMAX medium containing 1 μl each of Syn-GCaMP6f and TH-tdTomato virus (titer = 1 x 10^13^ genome copies/ml) was added to each dish and allowed to incubate at 37°C with 5% CO_2_ for 1 h, following which serum-free medium containing the AAVs were removed and replaced with 3 ml of supplemented neurobasal medium. Imaging was performed between 5 d after AAV infection.

### Imaging of spontaneous Ca^2+^ fluxes in cultured midbrain neurons

The protocol for imaging spontaneous Ca^2+^ fluxes in primary cultures of mouse midbrain neurons has been previously published (Bancroft & Srinivasan, 2020). Briefly, cultures were placed in a gas free recording buffer containing (mM): 154 NaCl, 5 KCl, 2 CaCl_2_, 0.5 MgCl_2_, 5 D-glucose, 10 HEPES, pH adjusted to 7.4 with NaOH (all purchased from Sigma-Aldrich). Imaging was performed using a confocal microscope (Fluoview 1200, Olympus) with a 40X, 0.8 NA water-immersion objective (Olympus). We used a 488-nm and 569-nm line of a krypton-argon laser to excite GCaMP6f and TH-tdTomato, respectively. The imaging frame was clipped to allow for a sampling rate of 1 frame per sec. For all experiments, spontaneous activity was imaged for 300 s, followed by peptide/drug application. Recording buffers were bath perfused using a peristaltic pump at a rate of 2 ml/min. For each field of view imaged a corresponding z-stack of both GCaMP6f and TH-tdTomato expression was captured and co-localization was used to identify TH^+^ cells. Diltiazem was purchased from Tocris (Minneapolis, MN), Mibefradil was purchased from Sigma-Aldrich, and cyclopiazonic acid (CPA) was purchased from Abcam.

### Data Analyses

Image processing was performed using ImageJ v1.52e (NIH, Rockville, MD, USA). To quantify S100B/TH ratios from immunostained mouse midbrain sections, z-stacks of TH labeled and corresponding S100B labeled midbrain sections were converted to projection images using the maximum intensity projection tool in ImageJ. TH and S100B labeled maximum intensity projections were manually thresholded and integrated fluorescence intensities from manually demarcated SNc and ventral tegmental area (VTA) midbrain regions were separately acquired for TH (red) and S100B (green) labels in each section. Integrated fluorescence intensity values for S100B and TH were used to derive ratios of S100B/TH for the SNc and VTA. S100B/TH ratios were used to measure S100B density in the SNc and VTA.

For analysis of Ca^2+^ flux kinetics in TH^+^ and TH^−^ neurons from primary mouse midbrain cultures, time stacks (t-stacks) obtained from confocal imaging of live midbrain neurons were used to create max intensity projections and adjusted to increase brightness in order to visualize all active neuronal somata in a field of view. Somata displaying spontaneous Ca^2+^ fluxes were individually traced with the polygon tool and added to the ROI manager. The ROIs were used to extract neuronal Ca^2+^ flux traces from t-stacks, and only somata with one or more spontaneous Ca^2+^ flux events were selected for analysis. Mean gray values from Ca^2+^ flux event traces were converted to ΔF/F values using a 10 s period with no Ca^2+^ flux events to obtain baseline fluorescence (F). For each XY time-series movie analyzed, a corresponding z-stack image of both hSyn-GCaMP and TH-tdTomato expression was used to create GCaMP and tdTomato co-localized max intensity projection images. Images obtained in this way were merged, and co-localization of GCaMP with tdTomato was used to definitively identify TH^+^ neurons, while cells displaying GCaMP labeling without tdTomato were identified as TH^−^ neurons. All neuronal Ca^2+^ flux traces obtained using the above procedures were manually analyzed with MiniAnalysis v6.0.7 (Synaptosoft, Fort Lee, NJ, USA) to detect and quantify Ca^2+^ flux frequency (fluxes/min) and amplitude (ΔF/F).

### Statistical analyses

All data and statistical analyses were performed using Origin 2019 v9.6 (OriginLab, Northampton, Massachusetts, USA). Data presented are mean ± s.e.m. For each data set, normality was first determined in Origin using the Shapiro-Wilk test. Normally distributed data were analyzed via two-sample t-tests or paired sample t-tests when appropriate. Non-normally distributed data were analyzed with Mann-Whitney or paired sample Wilcoxon-signed rank test for between group differences. Data was considered to be significantly different at *p* < 0.05. Statistical tests and sample sizes for each experiment are described in the figure legends, and exact *p* values comparing datasets are shown in the figures.

## Results

### The ratio of S100B expression to TH staining is higher in the SNc than the VTA

We first assessed the extent of S100B expression in the midbrain of adult mice. To do this, we imaged and quantified S100B expression in the SNc and VTA from midbrain sections of 3 – 4 month old adult male mice immunostained for TH and S100B (Fig. 1A). Interestingly, S100B containing astrocytic processes completely enveloped TH^+^ neuronal somata in the SNc, but not in the VTA (Fig. 1B). Quantification of fluorescence intensities for TH and S100B from multiple sections and mice showed that the VTA contained significantly higher levels of TH and S100B expression than the SNc (Fig. 1C). However, despite higher total expression of TH and S100B in the VTA, the ratio of S100B to TH intensity was ∼1.4 fold higher in the SNc than the VTA (Fig. 1D). Together, these data show that astrocytic S100B is robustly expressed in close proximity to the somata of SNc DA neurons, and that the SNc possesses a significantly higher ratio of S100B to TH expression when compared to the VTA.

**Figure 1.**
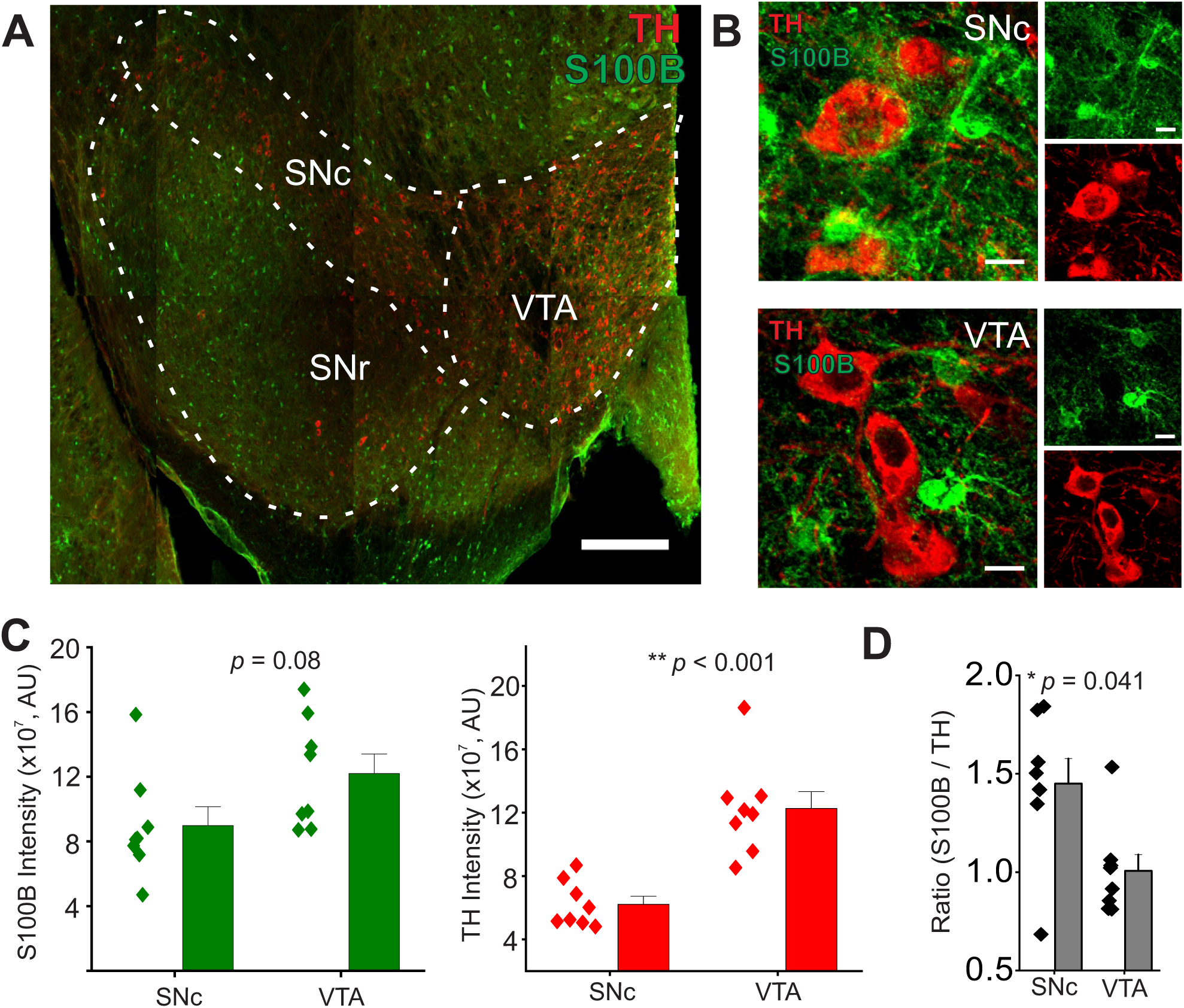
S100B density is higher in the SNc when compared to the VTA. **(A)** A representative confocal mosaic of a mouse midbrain section immunostained for S100B (green) and TH (red), scale bar = 300 μm. Subregions of the midbrain are indicated with dotted lines (SNc = substantia nigra pars compacta; VTA = ventral tegmental area; SNr = substantia nigra pars reticulata) **(B)** Representative high magnification confocal images of S100B and TH expression in the SNc (top) and VTA (bottom), scale bar = 10 μm. **(C)** The left graph shows the integrated S100B fluorescence intensity in the SNc and VTA. Right graph shows integrated TH fluorescence intensity in the SNc and VTA **(D)** Ratio of S100B/TH integrated intensity for the SNc and VTA. n = 8 midbrain sections from 3 male mice. All errors are SEM; p values are based on two sample t-tests.

### TH^+^ and TH^−^ neurons from primary mouse midbrain cultures show significant differences in the kinetics of spontaneous Ca^2+^ fluxes

Recent studies have shown that exposure to extracellular S100B alters the firing properties of pacemaking neurons (Morquette et al., 2015; Ryczko et al., 2021). Based on these findings, we rationalized that any abnormal increase in extracellular S100B concentrations in the SNc could similarly alter DA neuron function. To test our hypothesis, we cultured primary mouse ventral midbrain neurons from ED14 embryos, and maintained primary cultures for 19 days *in vitro* (DIV) prior to imaging (Fig. 2A) (Bancroft & Srinivasan, 2020). Immunostaining showed that midbrain cultures contain mature NeuN^+^ neurons that are either TH^+^ or TH^−^ (Fig. 2B) (Bancroft & Srinivasan, 2020; Srinivasan et al., 2016; Zarate et al., 2020). These neurons are co-cultured with primary midbrain astrocytes expressing S100B (Fig. 2B).

**Figure 2.**
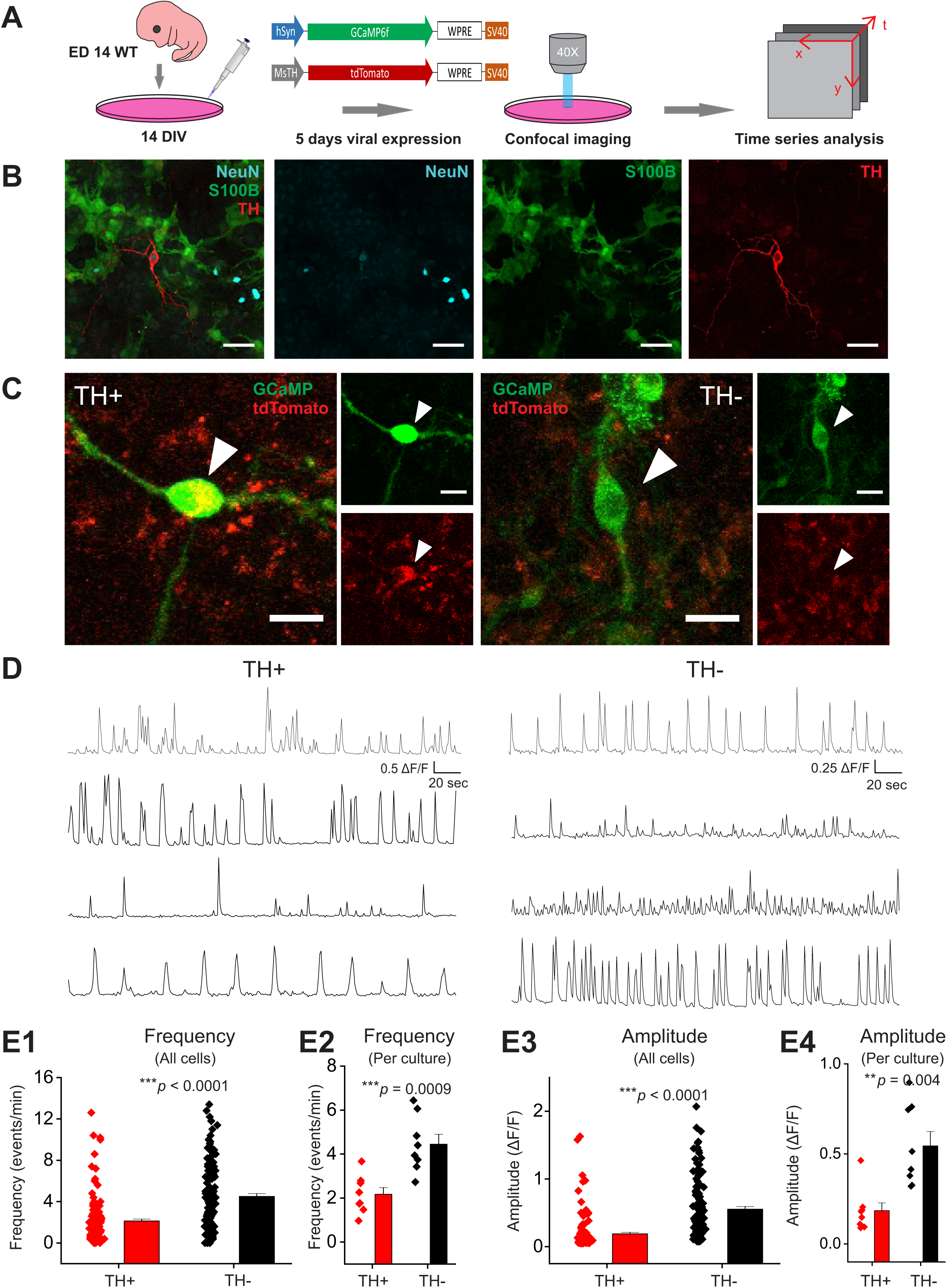
Cultured mouse primary TH^+^ and TH^−^ neurons differ with regard to spontaneous Ca^2+^ flux kinetics. **(A)** Schematic for measuring spontaneous Ca^2+^ fluxes in primary mouse midbrain neuron cultures using AAV 2/5 hSyn-GCaMP6f and AAV 2/5 TH-tdTomato viruses. **(B)** Representative image of a formalin-fixed primary mouse midbrain culture stained for NueN, S100B and TH; scale bar = 50 μm. **(C)** Representative confocal images of live primary mouse midbrain cultures are shown. Arrowheads point to an example of a TH^+^ neuron with hSyn-GCaMP6f (green) and TH-tdTomato (red) expression and an example of a TH^−^ neuron with hSyn-GCaMP6f expression, but no expression of TH-tdTomato; scale bar = 20 μm. **(D)** Multiple representative traces of spontaneous Ca^2+^ fluxes in TH^+^ and TH^−^ neurons showing the heterogenous nature of spontaneous Ca^2+^ flux events in cultured TH^+^ and TH^−^ neurons. **(E1)** Average frequency of Ca^2+^ flux events from individual TH^+^ and TH^−^ neurons **(E2)** Average frequency of TH^+^ and TH^−^ neurons binned by independent cultures **(E3)** Average amplitude of Ca^2+^ flux events from individual TH^+^ and TH^−^ neurons **(E4)** Average amplitude of TH^+^ and TH^−^ neurons binned by independent cultures. n = 137 neurons for TH^+^ cells and 134 neurons for TH^−^ cells from 8 independent weeks of culture. All errors are SEM; p values are based on Mann-Whitney tests.

To measure spontaneous Ca^2+^ fluxes in cultured midbrain neurons, we co-infected our cultures with AAV 2/5 hSyn-GCaMP6f and AAV 2/5 TH-tdTomato, which allowed the measurement of spontaneous Ca^2+^ fluxes in midbrain neurons, along with the ability to distinguish between TH^+^ and TH^−^ neurons. Five-day co-expression of GCaMP6f and tdTomato in midbrain cultures resulted in a robust expression of GCaMP6f in TH^+^ and TH^−^ neurons with tdTomato being expressed only in a subset of the total GCaMP6f expressing neuronal population (Fig. 2C). Since hSyn-GCaMP6f and TH-tdTomato expressing AAVs were co-applied to cultures at equivalent titers (1 x 10^9^ total genome copies per culture), and are both of the same serotype (AAV 2/5), the presence of tdTomato expression specifically reports TH^+^ neurons, while no tdTomato expression is indicative of a TH^−^ neuron (Fig. 2C).

Both TH^+^ and TH^−^ neurons exhibited robust spontaneous Ca^2+^ fluxes with heterogenous frequencies and amplitudes (Fig. 2D and supplementary movie 1) (Bancroft & Srinivasan, 2020). To control for the heterogenous nature of Ca^2+^ fluxes in these neurons, and the potential for skewing data with abnormal Ca^2+^ fluxes in one or more individual cultures, all TH^+^ and TH^−^ neurons were also binned by average frequencies and amplitudes obtained from all TH^+^ and TH^−^ neurons imaged per culture. We found that TH^−^ cells displayed a ∼2-fold higher Ca^2+^ flux frequency than TH^+^ neurons as well as a ∼2-fold higher amplitude in Ca^2+^ flux events when compared to TH^+^ neurons. The observed differences in neuronal Ca^2+^ flux frequency and amplitude persisted following binning, suggesting that the differences in Ca^2+^ flux kinetics between TH^+^ and TH^−^ neurons are due to inherent properties of TH^+^ and TH^−^ neurons rather than differences in Ca^2+^ fluxes between independent cultures (Fig. 2E1 – E4). These data show that cultured mouse midbrain neurons possess spontaneous Ca^2+^ fluxes, along with baseline differences between TH^+^ and TH^−^ neurons in spontaneous Ca^2+^ flux frequency and amplitude.

### Acute exposure to S100B increases spontaneous Ca^2+^ flux frequency only in TH^+^ neurons

Having observed spontaneous Ca^2+^ fluxes in cultured midbrain neurons, we next sought to determine if exposure to extracellular S100B alters spontaneous Ca^2+^ fluxes in TH^+^ and TH^−^ neurons. To do this, midbrain cultures were bath perfused with 50 pM of S100B peptide, and spontaneous Ca^2+^ flux frequencies and amplitudes in TH^+^ and TH^−^ neurons were measured. S100B application caused a significant 2-fold increase in the frequency of Ca^2+^ fluxes in TH^+^ neurons, but not in TH^−^ neurons (Figs. 3A and B, supplementary movie 1). The ability of extracellular S100B to specifically increase Ca^2+^ flux frequency only in TH^+^ neurons was also observed when data were plotted as average Ca^2+^ flux frequencies across multiple independent DA cultures (Fig. 3B), which rules out a skewing of the data as a result of a few individual DA cultures with abnormal S100B responses.

**Figure 3.**
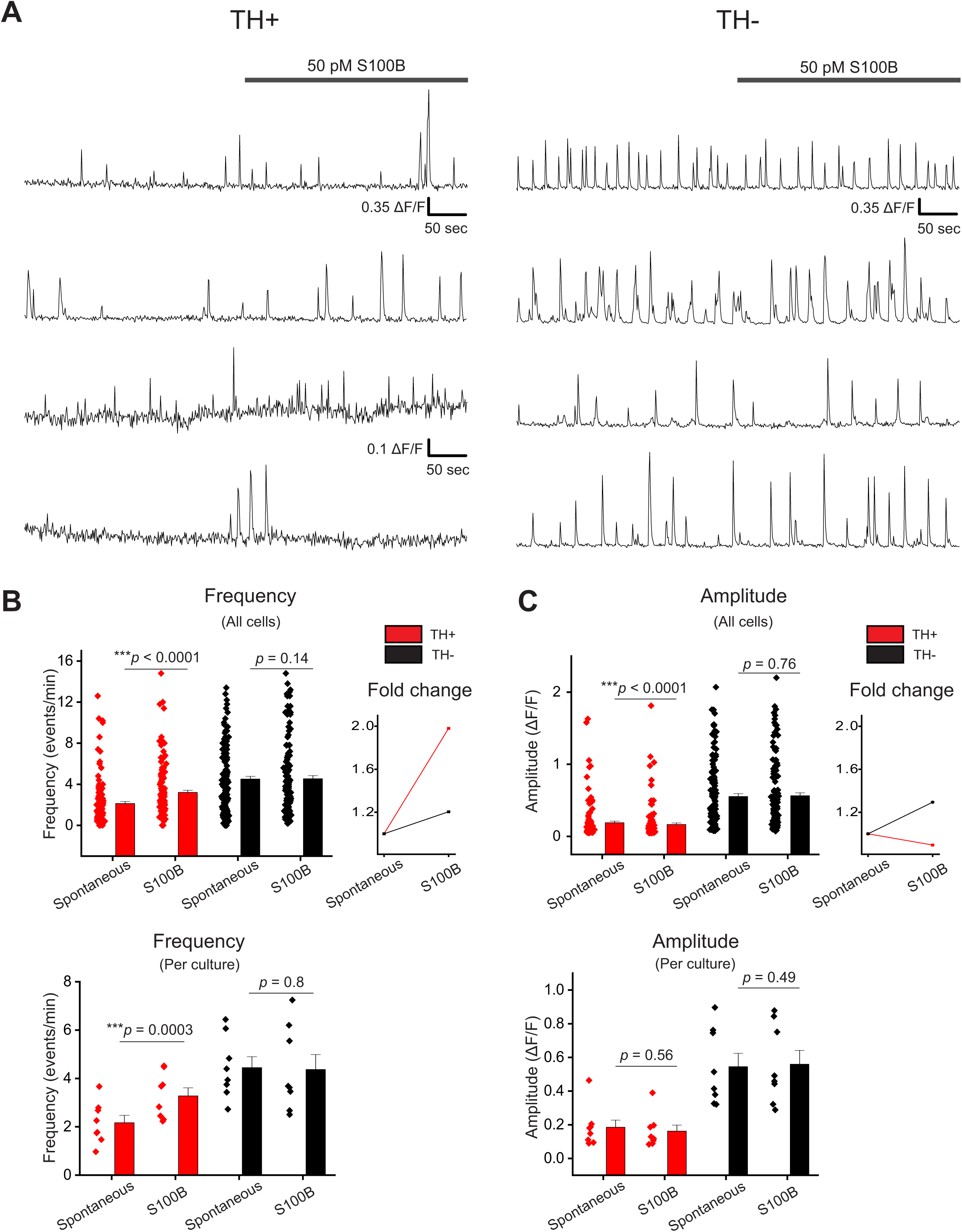
Acute exposure of primary midbrain cultures to S100B peptide increases spontaneous Ca^2+^ flux frequency only in TH^+^ neurons. **(A)** Multiple representative traces of spontaneous Ca^2+^ fluxes in TH^+^ and TH^−^ neurons with acute bath application of 50 pM S100B peptide are shown **(B)** A graph with average frequency of Ca^2+^ flux events from individual TH^+^ (red) and TH^−^ (black) neurons with and without S100B peptide is shown. The line graph on the right shows the average fold change of Ca^2+^ flux frequency for TH^+^ and TH^−^ cells with and without S100B. The graph below shows the average frequency of TH^+^ and TH^−^ neurons binned by week of culture **(C)** A graph with average amplitude of Ca^2+^ events from individual TH^+^ (red) and TH^−^ (black) neurons with and without S100B peptide. The line graph on the right shows the average fold change of Ca^2+^ flux amplitude for TH^+^ and TH^−^ cells with and without S100B. The graph below shows the average amplitude of TH^+^ and TH^−^ neurons binned by individual culture. n = 137 neurons per condition for TH^+^ cells and 134 neurons per condition for TH^−^ cells from 8 independent cultures. All errors are SEM; p values are based on Wilcoxon signed rank test.

S100B did not affect Ca^2+^ flux amplitudes in either TH^+^ or TH^−^ neurons when the data were plotted either as individual neurons or as average amplitudes from independent DA cultures (Fig. 3A and C), indicating that the effect of S100B on TH^+^ neurons occurred consistently across multiple independent cultures. It should be noted that we observed a very small, statistically significant reduction in Ca^2+^ flux amplitudes for TH^+^ neurons only when the data were plotted as an average of amplitudes from individual neurons and not across independent DA cultures.

Together, these data show that acute exposure to extracellular S100B specifically increases the frequency, but not the amplitude of spontaneous Ca^2+^ fluxes in TH^+^ neurons, with no effect on the frequency or amplitude of Ca^2+^ fluxes in TH^−^ neurons.

### Spontaneous Ca^2+^ fluxes in TH^+^ and TH^−^ neurons depend on extracellular Ca^2+^

The observation that extracellular S100B specifically increased Ca^2+^ flux frequency in TH^+^ neurons led us to assess sources of spontaneous Ca^2+^ fluxes in these neurons. Extracellular and intracellular Ca^2+^ stores were systematically depleted and neuronal Ca^2+^ fluxes were recorded under each condition. Exposure to zero Ca^2+^ aCSF caused a significant reduction in the frequency of Ca^2+^ fluxes in TH^+^ and TH^−^ neurons (Fig. 4A, supplementary movie 2). The few Ca^2+^ flux events that persisted in the presence of zero Ca^2+^ aCSF did not show any significant differences in amplitude for either TH^+^ or TH^−^ neurons (Fig. 4A).

**Figure 4.**
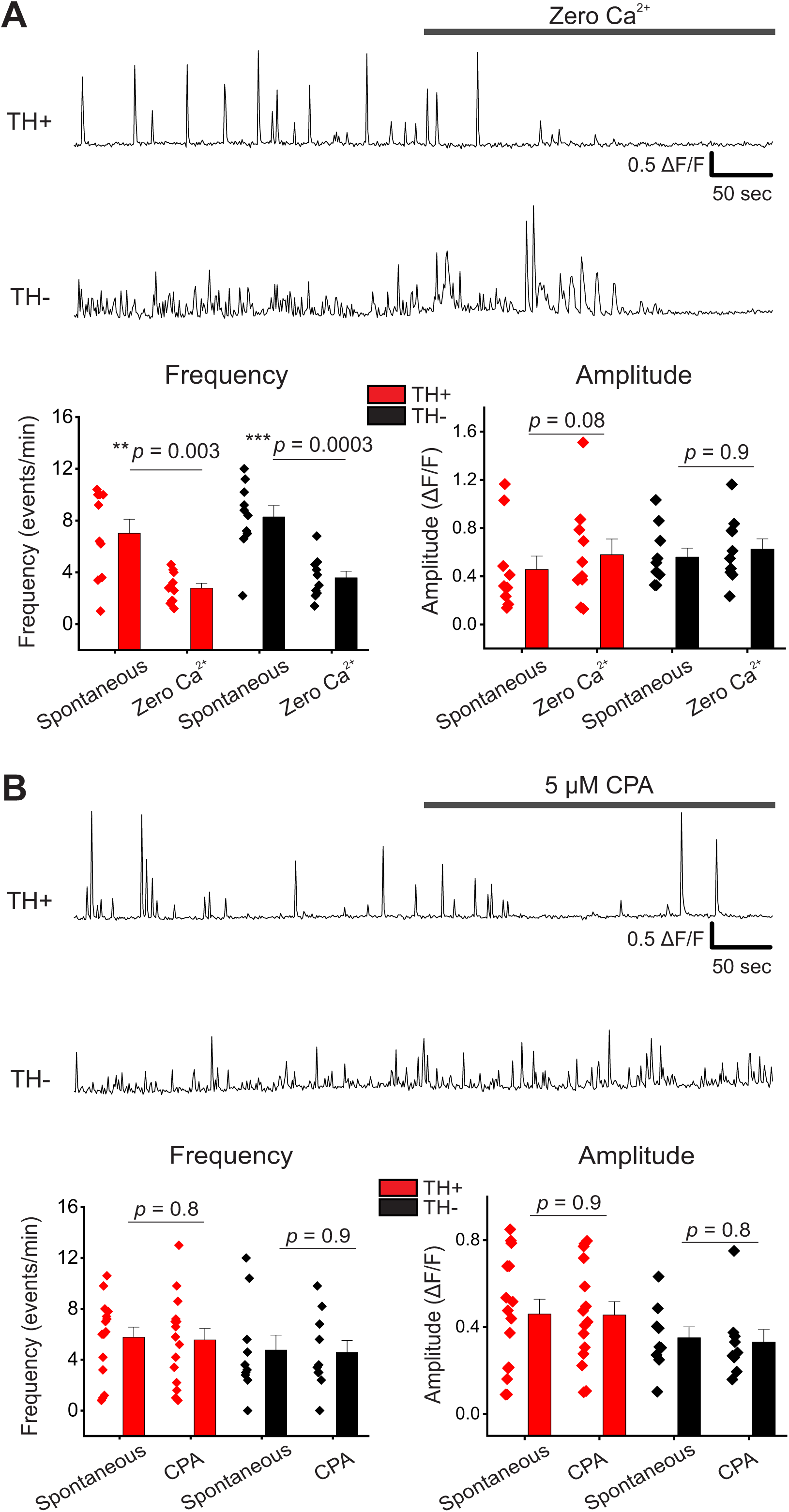
Spontaneous Ca^2+^ fluxes in cultured mouse midbrain neurons depend on extracellular Ca^2+^. **(A)** Representative traces of spontaneous Ca^2+^ fluxes in a TH^+^ and TH^−^ neuron with bath applied zero Ca^2+^ aCSF. The graphs below show average Ca^2+^ flux frequency and amplitude for TH^+^ (red) and TH^−^ (black) neurons with and without zero Ca^2+^ aCSF **(B)** Representative traces of spontaneous Ca^2+^ fluxes in a TH^+^ and TH^−^ neuron with bath applied 5 μM CPA. The graphs below show average Ca^2+^ flux frequency and amplitude for TH^+^ (red) and TH^−^ (black) neurons with and without 5 μM CPA. For zero Ca^2+^ aCSF experiments, n = 10 neurons for TH^+^ cells and n = 15 neurons for TH^−^ cells from 3 independent weeks of culture. For CPA experiments, n = 15 neurons for TH^+^ cells and 10 neurons for TH^−^ cells from 3 independent weeks of culture. All errors are SEM; p values are based on paired sample t tests or Wilcoxon signed ranked tests as appropriate.

To assess the extent to which intracellular Ca^2+^ contributes to neuronal Ca^2+^ fluxes, endoplasmic reticulum (ER) Ca^2+^ stores were depleted with a 15 min exposure to 5 μM cyclopiazonic acid (CPA), which is an inhibitor of the Ca^2+^-ATPase SERCA pump in the ER. CPA did not affect spontaneous Ca^2+^ flux frequency or amplitude in either TH^+^ or TH^−^ neurons (Fig. 4B, supplementary movie 3). Thus, spontaneous Ca^2+^ fluxes in primary mouse DA cultures primarily depend on extracellular Ca^2+^, with very little dependence on ER Ca^2+^ stores.

### L-type VGCCs largely contribute to spontaneous Ca^2+^ fluxes in TH^+^ and TH^−^ neurons

Because our data showed that spontaneous Ca^2+^ fluxes in cultured midbrain neurons depend primarily on extracellular Ca^2+^ (Fig. 4A), and since midbrain neurons require L-type VGCC activity for pacemaking (Chan et al., 2007; Guzman, Sanchez-Padilla, Chan, & Surmeier, 2009; Leandrou, Emmanouilidou, & Vekrellis, 2019; Puopolo, Raviola, & Bean, 2007; Putzier, Kullmann, Horn, & Levitan, 2009), we assessed the extent to which L-type VGCCs contribute to spontaneous Ca^2+^ fluxes in TH^+^ and TH^−^ neurons. Ca^2+^ fluxes were recorded in TH^+^ and TH^−^ neurons following acute exposure of midbrain cultures to either the L-type VGCC blocker, diltiazem or the T-type VGCC blocker, mibefradil. 100 µM diltiazem significantly decreased the frequency of spontaneous Ca^2+^ fluxes in TH^+^ and TH^−^ neurons. Following exposure to diltiazem, Ca^2+^ flux frequency was reduced by ∼5-fold in TH^+^ neurons and 2.5 fold in TH^−^ neurons (Fig. 5A). Interestingly, only TH^+^ neurons displayed a significant, 1.5-fold change in Ca^2+^ flux amplitude in Ca^2+^ fluxes that persisted in the presence of diltiazem (Fig. 5A).

**Figure 5.**
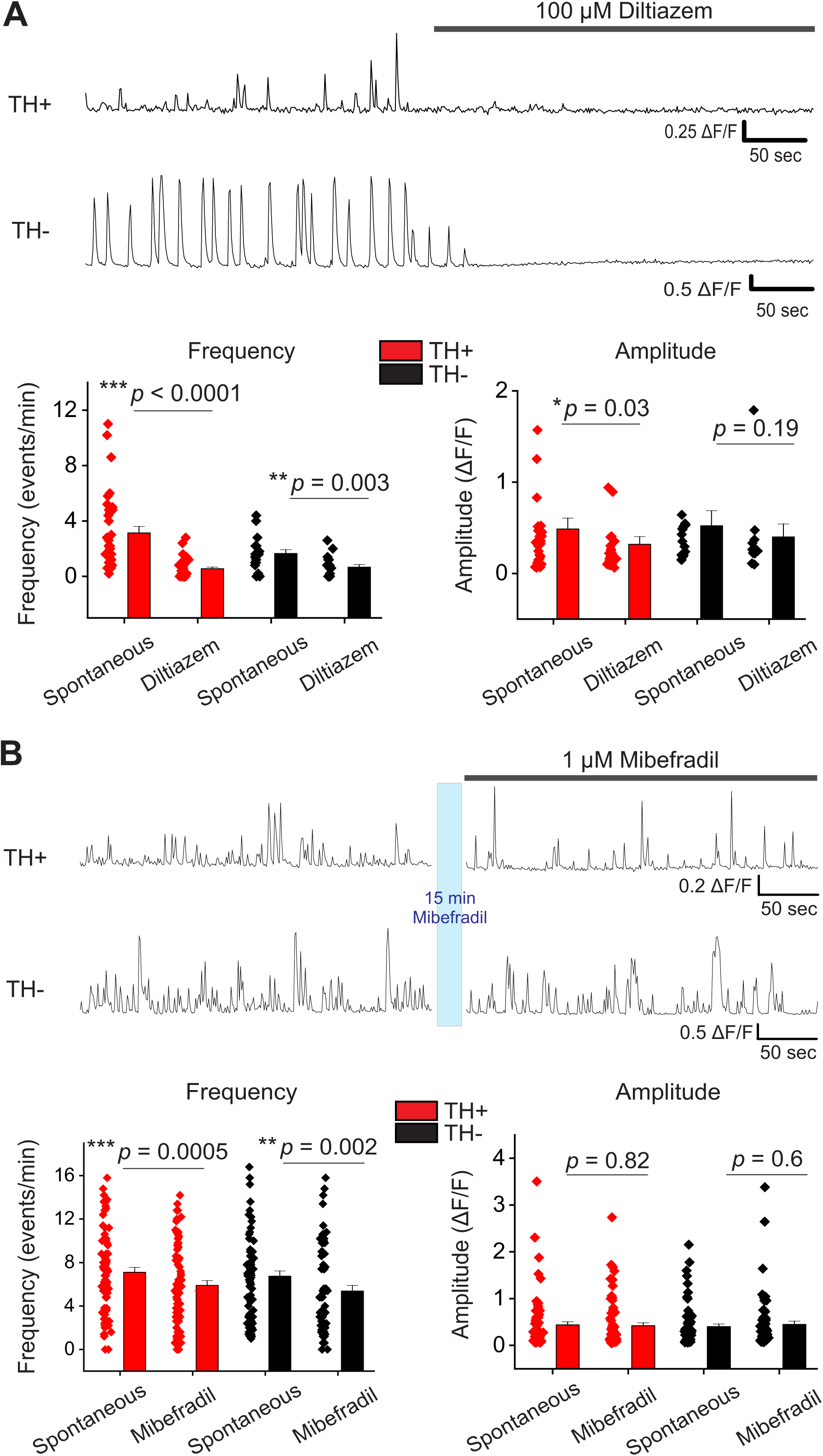
Spontaneous Ca^2+^ fluxes in cultured mouse midbrain neurons largely require L-type VGCCs. **(A)** Representative traces of spontaneous Ca^2+^ fluxes in a TH^+^ and TH^−^ neuron with bath applied 100 μM diltiazem. The graphs below show average Ca^2+^ flux frequency and amplitude of TH^+^ (red) and TH^−^ (black) neurons with and without diltiazem **(B)** Representative traces of spontaneous Ca^2+^ fluxes in a TH^+^ and TH^−^ neuron following 15 min bath application of 1 μM mibefradil. The graphs below show average Ca^2+^ flux frequency and amplitude of TH^+^ (red) and TH^−^ (black) neurons with and without mibefradil. For diltiazem experiments, n = 34 neurons for TH^+^ cells and 18 neurons for TH^−^ cells from 4 independent weeks of culture. For mibefradil experiments, n = 75 neurons for TH^+^ cells and 66 neurons for TH^−^ cells from 4 independent weeks of culture. All errors are SEM; p values are based on Wilcoxon signed rank test.

To assess the contribution of T-type VGCCs to Ca^2+^ fluxes, midbrain cultures were exposed to 1 µM mibefradil. Mibefradil decreased Ca^2+^ flux frequencies by ∼1.2-fold for both TH+ and TH-neurons (Fig. 5B, supplementary movie 5). These data show that a majority of spontaneous Ca^2+^ fluxes in cultured midbrain neurons occur due to the activation of L-type VGCCs and that L-type VGCCs are more active in TH^+^ than in TH^−^ neurons.

### L-type VGCCs are necessary for S100B-mediated increases in the spontaneous Ca^2+^ flux frequency of TH^+^ neurons

Having found that a majority of spontaneous Ca^2+^ fluxes in TH^+^ neurons are mediated by L-type VGCCs, we asked if S100B requires L-type VGCCs for increasing Ca^2+^ flux frequency in TH^+^ neurons. Spontaneous Ca^2+^ fluxes in midbrain neurons were recorded following bath application of 50 pM S100B, and a subsequent co-application of 50 pM S100B with 100 µM diltiazem (Fig. 6A). Acute application of S100B significantly increased Ca^2+^ flux frequency in TH^+^ neurons, but not TH^−^ neurons and did not alter the amplitude of Ca^2+^ fluxes in either TH^+^ or TH^−^ neurons (Fig. 6B and C). Co-exposure of midbrain cultures to 50 pM S100B + 100 μM diltiazem completely inhibited the S100B-mediated increase of Ca^2+^ fluxes in TH^+^ neurons, with no effect on Ca^2+^ flux amplitude for the few remaining neuronal Ca^2+^ events (Fig. 6B and C, supplementary movie 4). These results clearly demonstrate that S100B increases Ca^2+^ flux frequency in TH^+^ neurons via L-Type VGCCs.

**Figure 6.**
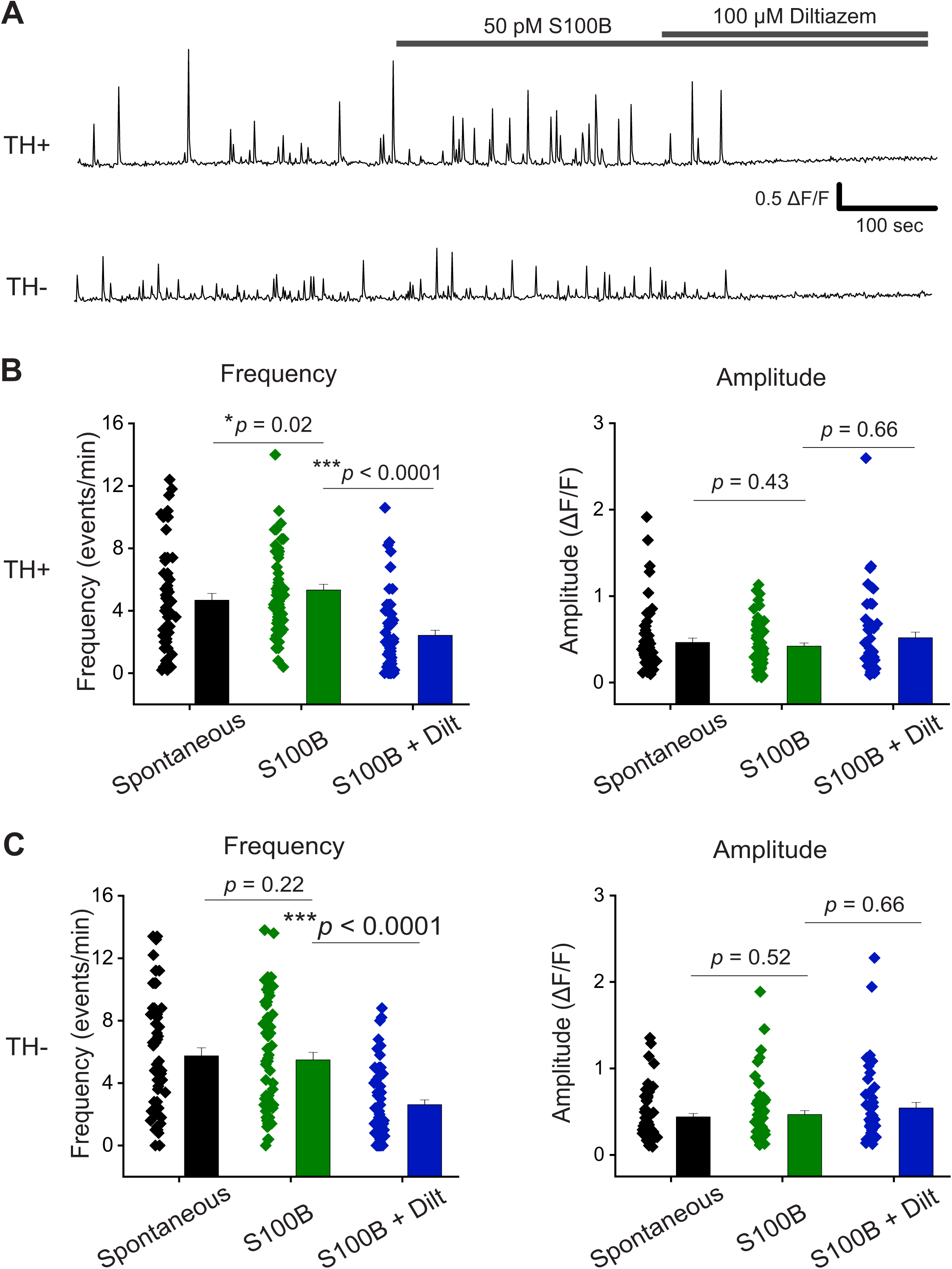
Extracellular S100B increases spontaneous Ca^2+^ fluxes in TH^+^ DA neurons via L-type VGCCs. **(A)** Representative traces of spontaneous Ca^2+^ fluxes in a TH^+^ and TH^−^ neuron with bath application of S100B, followed by co-application of S100B with diltiazem **(B)** Graphs show average Ca^2+^ flux frequency and amplitude of TH^+^ neurons without any drug (black) with bath applied S100B (green), and co-applied S100B + diltiazem (blue) **(C)** Graphs show average Ca^2+^ flux frequency and amplitude of TH^−^ neurons without drug (black) with bath applied S100B (green), and co-applied S100B + diltiazem (blue). n = 55 neurons for TH^+^ cells and 57 neurons for TH^−^ cells from 4 independent weeks of culture. All errors are SEM; p values are based on Wilcoxon signed rank test.

### T-type VGCCs are not involved in the S100B-mediated increase of Ca^2+^ flux frequencies in TH^+^ neurons

To determine if T-type VGCCs are involved in S100B-medaited increases in the Ca^2+^ flux frequency of TH^+^ DA neurons, spontaneous Ca^2+^ fluxes were recorded in aCSF, then treated for 15 min with the T-type VGCC antagonist, mibefradil. Ca^2+^ fluxes in TH^−^ and TH^+^ DA neurons were then recorded in the presence of 1 µM mibefradil + 50 pM S100B (Fig. 7A). We found that mibefradil significantly reduced Ca^2+^ flux frequency in both TH^+^ and TH^−^ neurons (Fig. 7B and C). However, co-application of S100B with mibefradil was unable to inhibit S100B-mediated increases in Ca^2+^ flux frequency in TH^+^ DA neurons (Fig. 7B, supplementary movie 5). In addition, amplitudes for TH^+^ neurons were mostly unchanged across all recording conditions (Fig. 7B). In the case of TH^−^ neurons, we observed significant reductions in Ca^2+^ flux frequency following exposure to either mibefradil alone or co-exposure of mibefradil with S100B (Fig. 7C) In addition, none of the recording conditions altered Ca^2+^ flux amplitudes in TH^+^ and TH^−^ neurons (Fig. 7C). Together, these results suggest that the effect of S100B-mediated increases in Ca^2+^ flux frequency in TH^+^ neurons does not depend on T-type VGCCs.

**Figure 7.**
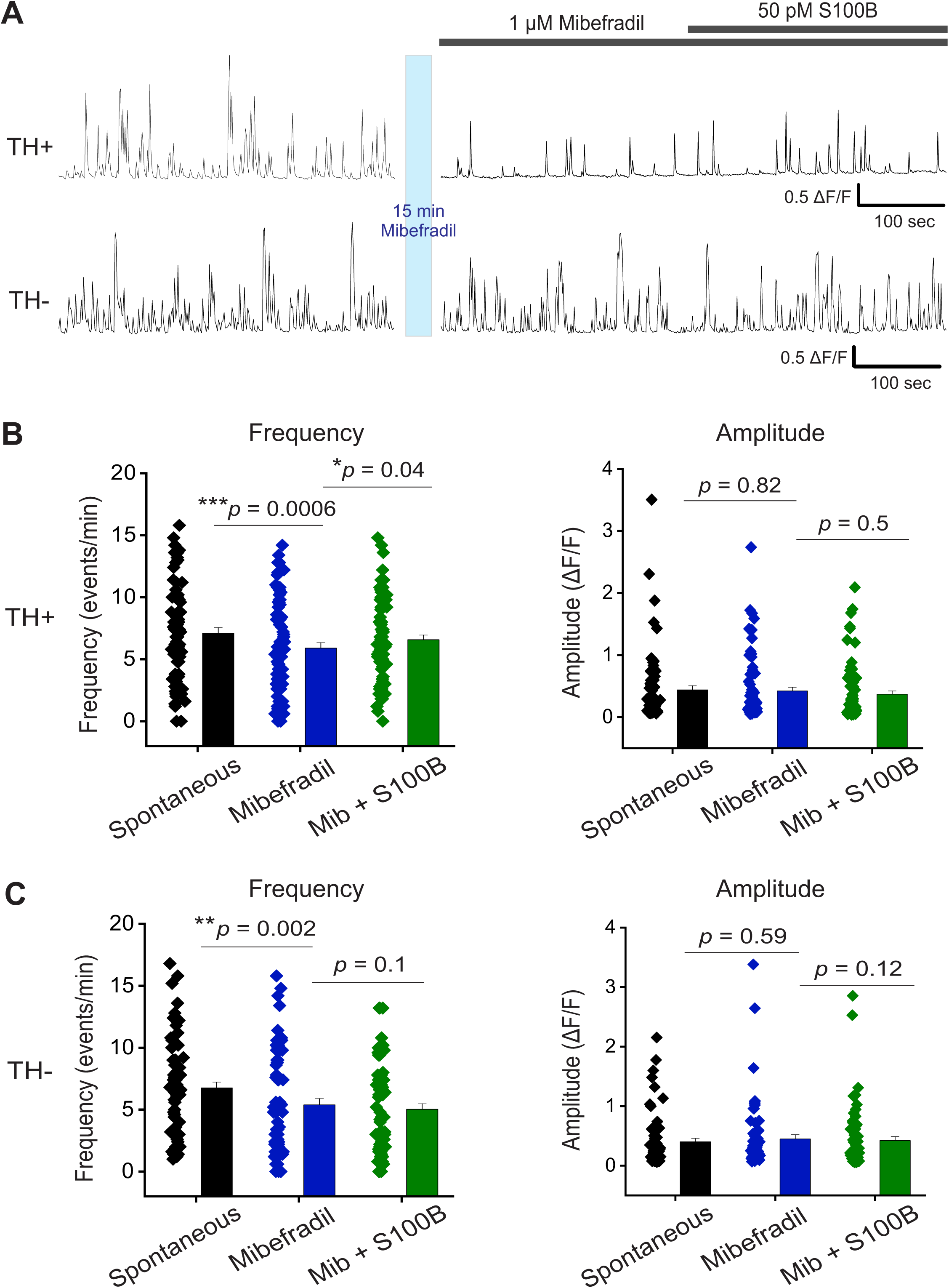
T-type VGCCs are not required for S100B-mediated increases of spontaneous Ca^2+^ fluxes in TH^+^ DA neurons. **(A)** Representative traces of spontaneous Ca^2+^ fluxes in a TH^+^ and TH^−^ neuron after 15 min incubation with 1 μM mibefradil, followed by co-application of S100B with mibefradil **(B)** Graphs show average Ca^2+^ flux frequency and amplitude of TH^+^ neurons without any drug (black) with bath applied mibefradil (blue), and co-applied S100B + mibefradil (green) **(C)** Graphs show average Ca^2+^ flux frequency and amplitude of TH^−^ neurons without drug (black) with bath applied mibefradil (blue), and co-applied S100B + mibefradil (green). n = 75 neurons for TH^+^ cells and 66 neurons for TH^−^ cells from 4 independent weeks of culture. All errors are SEM; p values are based on Wilcoxon signed rank test.

### Chronic S100B differentially alters the kinetics of Ca^2+^ fluxes in TH^+^ and TH-neurons

Since acute exposure to S100B specifically increased the frequency of Ca^2+^ events only in TH^+^ neurons, we assessed the extent to which chronic extracellular exposure to S100B alters spontaneous Ca^2+^ events in cultured midbrain neurons. 50 pM of the S100B peptide was added to the neuronal media and incubated for 7 days prior to Ca^2+^ imaging. We found that chronic S100B had distinct effects on the Ca^2+^ flux kinetics of TH^+^ and TH^−^ neurons (Fig. 8A, supplementary movie 6).

**Figure 8.**
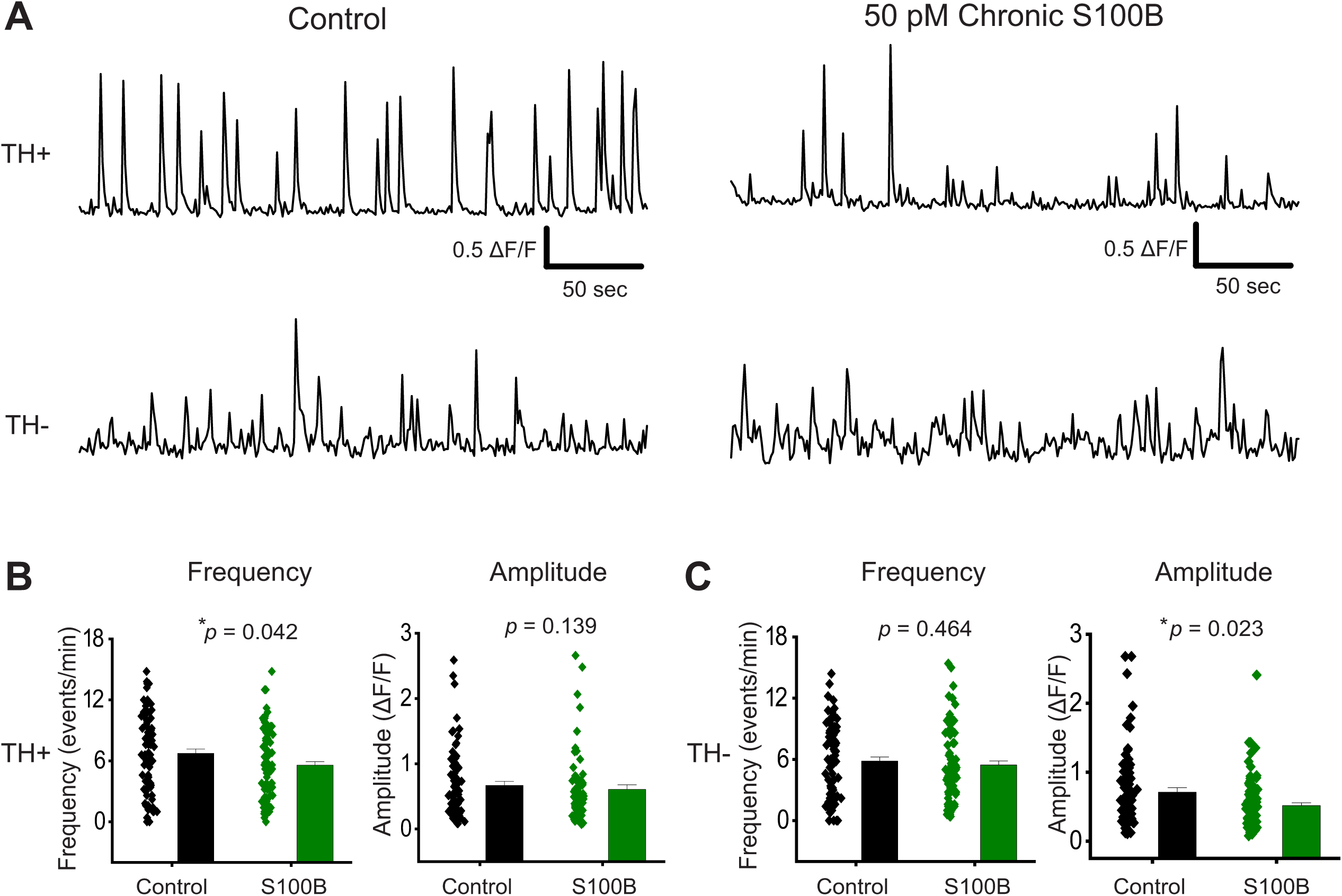
Chronic S100B reduces the frequency of Ca^2+^ fluxes in TH^+^ neurons and the amplitude of Ca^2+^ fluxes in TH^−^ neurons. **(A)** Representative traces of spontaneous Ca^2+^ fluxes in TH^+^ and TH^−^ neurons without any treatment (Control, left traces), and following a 7-day chronic incubation with 50 pM S100B (right traces) **(B)** Average frequency and amplitude of spontaneous Ca^2+^ events of TH^+^ neurons in untreated control (black) and chronic S100B (green) treated conditions **(C)** Average frequency and amplitude of spontaneous Ca^2+^ events of TH^−^ neurons in untreated control (black) and chronic S100B (green) treated conditions. For control condition, n = 78 neurons for TH^+^ cells and 79 neurons for TH^−^ cells from 4 independent weeks of culture. For chronic S100B condition, n = 89 neurons for TH^+^ cells and 86 neurons for TH^−^ cells from 4 independent weeks of culture. All errors are SEM; p values are based on Mann-Whitney tests.

In the case of TH^+^ neurons, S100B exposure caused a significant 3-fold reduction in spontaneous Ca^2+^ flux frequencies when compared to untreated controls, while Ca^2+^ event amplitudes for both conditions were similar (Fig. 8B). By contrast, chronic S100B significantly reduced Ca^2+^ flux amplitudes in TH^−^ neurons by ∼4-fold when compared to untreated controls, while Ca^2+^ event frequencies in TH^−^ neurons remained unaffected (Fig. 8C). Together, these results demonstrate that chronic exposure to extracellular S100B dramatically alters the kinetics of spontaneous Ca^2+^ events in TH^+^ and TH^−^ midbrain neurons, albeit in different ways.

## Discussion

Developing effective neuroprotective therapies for PD requires an understanding of the specific pathological mechanisms leading to DA neuron loss in the SNc. In this context, the glial-specific protein, S100B is of particular interest because multiple reports in humans and in animal models of PD suggest that S100B plays an active role in the degeneration of SNc DA neurons (D. Z. Carvalho et al., 2015; Fardell et al., 2018; Hohoff et al., 2010; Liu et al., 2011; Sathe et al., 2012; Schaf et al., 2005). Based on these reasons, our study focuses on how exposure to extracellular S100B alters the activity of DA neurons.

We show that S100B-containing astrocytic processes envelop DA neuronal somata in the SNc, but not in the VTA (Fig. 1B). We also demonstrate an increased ratio of astrocytic S100B to TH staining in the SNc compared to the VTA (Fig. 1D). These findings are significant because astrocytic coverage of neuropil is a critical determinant in modulating the excitability of neurons via mechanisms such as the secretion of astrocyte-derived factors, the clearance of neurotransmitters, and extracellular K^+^ buffering (Verhoog, Holtman, Aronica, & van Vliet, 2020). The observed morphological differences in interaction of astrocytic processes with SNc DA neurons in combination with an increased S100B to TH expression ratio in the SNc when compared to the VTA, suggest that any abnormal increase in astrocytic secretion of S100B in the SNc could significantly alter the function of SNc DA neurons with relatively little effect on DA neurons within the VTA.

Recent studies show that secreted S100B can buffer extracellular Ca^2+^, thereby altering the function of Ca^2+^ sensitive K^+^ channels in pacemaking neurons of the trigeminal ganglion and the cortex (Morquette et al., 2015; Ryczko et al., 2021). Given the precedence for a mechanism by which extracellular S100B can modulate neuronal function in multiple brain regions, we sought to determine the extent to which extracellular S100B exposure alters the activity of DA neurons. In this study, we show for the first time that extracellular exposure to 50 pM S100B increases the frequency of spontaneous Ca^2+^ fluxes only in TH^+^ DA neurons by ∼2-fold (Fig. 3B). Since 50 pM of S100B cannot buffer extracellular Ca^2+^ to the extent of modulating Ca^2+^ sensitive K^+^ channels, our data suggest that S100B alters VGCCs governing spontaneous Ca^2+^ fluxes in DA neurons. This view is supported by our finding that inhibition of L-type, but not T-type VGCCs in DA neurons is sufficient to abolish the effect of extracellular S100B-mediated increases in Ca^2+^ flux frequency (Figs. 6 and 7). The average concentration of S100B levels in the CSF of PD is ∼3.1 μg/l (Sathe et al., 2012), and studies also show that ∼70 to 80% of S100B in the CSF is due to secretion from the brain and not from serum (Begcevic, Brinc, Drabovich, Batruch, & Diamandis, 2016; Sathe et al., 2012). Based on these findings, we estimate that 50 – 100 pM of extracellular S100B in the immediate vicinity of neurons would be an abnormal extracellular concentration of this protein in the midbrain during early stage PD. Thus, the fact that 50 pM extracellular S100B exposure specifically alters Ca^2+^ flux frequency in DA neurons is highly relevant to reported concentrations of S100B in the CSF of patients with clinical PD.

Multiple studies showing the central role of VGCCs in generating pacemaking activity in DA neurons (Chan et al., 2007; Guzman et al., 2009; Leandrou et al., 2019; Puopolo et al., 2007; Putzier et al., 2009) strongly indicate that VGCC-mediated Ca^2+^ fluxes are necessary for the generation of spontaneous action potentials in DA neurons. Our finding that spontaneous VGCC-mediated Ca^2+^ flux frequencies are significantly higher in TH^−^ neurons when compared to TH^+^ cells (Fig. 2) mirrors previously reported differences in spontaneous action potential firing between these two cell types in culture, with action potentials in TH^+^ neurons at 1 – 4 Hz, and TH^−^ neurons at ∼10 Hz (Srinivasan et al., 2016). Therefore, quantifying spontaneous VGCC-mediated Ca^2+^ fluxes with AAV-expressed genetically encoded calcium indicators is a reliable measure of the pacemaking activity in cultured midbrain neurons. Since VGCCs are critical for the generation of action potentials in pacemaking neurons, a small pathological disruption in either the frequency or magnitude of VGCC-mediated Ca^2+^ fluxes can have exponentially damaging consequences for action potential generation and dopamine release at DA neuron terminals. One can therefore surmise that the observed ∼2-fold increase in the frequency of L-type VGCC-mediated Ca^2+^ fluxes induced by extracellular S100B (Fig. 3) would completely alter somatodendritic, as well as axonal release of dopamine by SNc DA neurons, leading to an early pathological dysfunction in feedback control of basal ganglia circuitry. Another point to note is that an increase in extracellular S100B-mediated increases in the frequency rather than the amplitude of Ca^2+^ fluxes suggests that extracellular S100B likely increases the activation of dormant L-type VGCCs, resulting in an increased probability of synchronous L-type VGCC activation in DA neurons. Therefore, rather than being a direct agonist of VGCCs, S100B is likely an allosteric modulator of L-type VGCCs.

We found that extracellular S100B caused an increase in Ca^2+^ fluxes only in TH^+^ neurons with no effect on TH^−^ neurons (Fig. 3B). This finding suggests that TH^+^ neurons may possess a functionally distinct isoform of L-type VGCCs, and it is possible that DA neurons express a novel L-type VGCC isoform that enables specific interaction with extracellular S100B and consequently, the modulation of VGCC activity only in midbrain DA neurons. An alternative mechanism for the specific effect of extracellular S100B on TH^+^ neurons comes from the fact that distinct L-type VGCC isoforms can be modulated by G-protein coupled receptors (GPCRs) (Altier, 2012). However, our data show that the majority of spontaneous Ca^2+^ fluxes in DA neurons require an extracellular Ca^2+^ source (Fig. 4), and that the L-type VGCC inhibitor diltiazem prevents extracellular S100B-mediated increases in the frequency of Ca^2+^ fluxes in TH^+^ DA neurons (Fig. 6). The likelihood that extracellular S100B directly interacts with L-type VGCCs in DA neurons is further bolstered by that fact that S100B and the S100 family of proteins are intrinsically disordered (S. B. Carvalho et al., 2013), and known to interact with multiple ion channels (Hermann et al., 2012). Therefore, S100B may have a number of novel, as yet undiscovered extracellular targets, including novel variants of L-type VGCCs specifically expressed in DA neurons. In addition, our findings point to the idea that TH^+^ DA neurons may express an isoform of L-type VGCCs with a distinct pharmacologic and functional profile from L-type VGCCs in TH^−^ midbrain neurons.

Chronic exposure to extracellular S100B peptide caused a decrease in the Ca^2+^ flux frequency of TH^+^ DA neurons, while simultaneously decreasing Ca^2+^ flux amplitude in TH^−^ neurons (Fig. 8). We postulate at least two mechanistic interpretations for this finding. The first is a cell autonomous mechanism in which prolonged stimulation of L-type VGCCs expressed by TH^+^ neurons can cause a compensatory internalization of functional L-type VGCC channels from the surface of TH^+^ DA neurons, as has been previously shown for L-type VGCCs following increased electrical activity (Green, Barrett, Bultynck, Shamah, & Dolmetsch, 2007). This can result in the observed reduction in frequency of L-type VGCC-mediated spontaneous Ca^2+^ fluxes in TH^+^ neurons following chronic exposure to extracellular S100B. The second is a circuit-based mechanism in which chronic exposure to extracellular S100B can cause an abnormal increase in the somatodendritic release of dopamine from TH^+^ neurons, resulting in the activation of D1-like receptors in TH^−^ GABAergic neurons and a consequent increase in the amplitude of L-type VGCC-mediated spontaneous Ca^2+^ fluxes within TH^−^ neurons (Zhou, Jin, Matta, Xu, & Zhou, 2009). Based on these mechanistic interpretations, we infer that chronic extracellular exposure to picomolar concentrations of S100B can fundamentally, and pathologically alter the activity of SNc DA neurons, as well as substantia nigra pars reticulata (SNr) GABAergic neurons.

In conclusion, our novel finding that picomolar extracellular concentrations of S100B can specifically alter Ca^2+^ fluxes via L-type VGCC expressed in TH^+^ DA neurons is relevant for understanding the mechanisms involved during early stages of PD pathogenesis. In future studies, we will: (**i**) Assess the effect of extracellularly secreted S100B from SNc astrocytes on DA function, neuronal loss, and PD-related behaviors *in vivo*, (**ii**) Elucidate the molecular mechanism by which extracellular S100B specifically alters the function of L-type VGCCs in SNc DA neurons, and (**iii**) Develop strategies to disrupt pathological interactions between extracellular S100B and L-type VGCCs in SNc DA neurons as a translationally relevant neuroprotective approach for the treatment of early stage PD.

## Acknowledgements

We thank the Texas A&M Institute for Genomic Medicine (TIGM) for providing timed pregnant mice to culture midbrain neurons. This work was partially funded by a research grant from the American Parkinson Disease Association (APDA) to RS and a National Institutes of Health (NIH) research grant, R01NS115809 to RS

## Supplementary Movie Captions

**Supplementary Movie 1.** Ca^2+^ fluxes in a TH^+^ (red arrow) and TH^−^ (cyan arrow) neuron with 50 pM S100B

**Supplementary Movie 2.** Reduced Ca^2+^ fluxes with zero Ca^2+^ aCSF

**Supplementary Movie 3.** No effect on Ca^2+^ fluxes with CPA

**Supplementary Movie 4.** Diltiazem blocks S100B-mediated increases in Ca^2+^ fluxes. TH^+^ neuron (red arrow) and TH^−^ neuron (cyan arrow)

**Supplementary Movie 5.** Mibefradil fails to block S100B-mediated increases in Ca^2+^ fluxes. TH^+^ neuron (red arrow) and TH^−^ neuron (cyan arrow)

**Supplementary Movie 6.** Chronic S100B decreases Ca^2+^ flux frequency in TH^+^ neuron (red arrows) and increases Ca^2+^ amplitude in TH^−^ neuron (cyan arrow)

## Notes

**Conflict of interest statement:** Authors declare no competing financial interests

### Competing Interest Statement

The authors have declared no competing interest.

